# Knockout of zebrafish desmin genes does not cause skeletal muscle degeneration but alters calcium flux

**DOI:** 10.1101/2020.10.16.342485

**Authors:** Gulsum Kayman Kurekci, Ecem Kural Mangit, Cansu Koyunlar, Seyda Unsal, Berk Saglam, Bora Ergin, Merve Gizer, Ismail Uyanik, Niloufar Boustanabadimaralan Düz, Petek Korkusuz, Beril Talim, Nuhan Purali, Simon M. Hughes, Pervin R. Dincer

## Abstract

Desmin is a muscle-specific intermediate filament protein that has fundamental role in muscle structure and force transmission. Whereas human desmin protein is encoded by a single gene, two desmin paralogs (*desma* and *desmb*) exist in zebrafish. *Desma* and *desmb* show differential spatiotemporal expression during zebrafish embryonic and larval development, being similarly expressed in skeletal muscle until hatching, after which expression of *desmb* shifts to gut smooth muscle. We generated knockout (KO) mutant lines carrying loss-of-function mutations for each gene by using CRISPR/Cas9. Mutants are viable and fertile, and lack obvious skeletal muscle, heart or intestinal defects. In contrast to morphants, knockout of each gene did not cause any overt muscular phenotype, but did alter calcium flux in myofibres. These results point to a possible compensation mechanism in these mutant lines generated by targeting nonsense mutations to the first coding exon.

## Introduction

Desmin is a type III intermediate filament protein that is specifically expressed in skeletal, cardiac and smooth muscles. In addition to their fundamental role in maintaining the structural integrity of the sarcomere, desmin intermediate filaments are involved in mechanotransduction and organelle positioning. Desminopathies belonging to myofibrillar myopathies are primarily characterized by abnormal protein aggregates in muscle and present as progressive skeletal myopathy and/or cardiomyopathy^1^. Although the involvement of smooth muscles is not widely reported, some patients suffer from smooth muscle dysfunction such as swallowing difficulties, intestinal pseudo-obstruction and respiratory insufficiency^2–4^. The majority of desmin mutations are associated with desmin aggregates; however, some mutations have been reported to not alter filament assembly and network integrity *in vivo* or *in vitro*^5,6^. KO of desmin in mice does not affect the viability and the development of muscles; however, muscle degeneration and cardiomyopathy are observed^7,8^.

Zebrafish is a widely used model for neuromuscular disorders^9,10^. Besides the advantage of being the most abundant tissue in zebrafish; the gene profile, and the structural and histological features of mammalian skeletal muscle are highly preserved in zebrafish^11^. To date, two approaches have been used to study the effect of the loss of desmin in zebrafish. On one hand, it was shown that morpholino-mediated knockdown of desmin causes skeletal and cardiac muscle myopathy^12,13^. On the other hand, Ramspacher et al. studied a desmin mutant line (*desma*^*sa5*^), generated by ENU mutagenesis, which causes a truncation mutation approximately half way along the molecule, that also presented with skeletal and cardiac muscle phenotype^14^.

Gene duplications are common in zebrafish and most duplicated genes have similar functions to their human orthologs. Alternatively, paralogs can be expressed in different tissues or stages of development and have distinct functions to their human orthologs. Humans carry a single copy of the *DES* gene whereas two desmin genes (*desma* and *desmb*) are found in zebrafish^13^. Previous work has not established the tissue specific distribution and role of these two desmin paralogs.

Here, we report the spatiotemporal expression pattern of the two desmin paralogs in zebrafish, as well as the generation using CRISPR/Cas9 and characterization of putative null zebrafish mutant lines for each of the desmin paralogs. We show that *desma* and *desmb* are expressed in skeletal muscle until 72-hour post-fertilization (hpf). After 72 hpf, expression of *desmb* shifts to gut smooth muscle. Putative null mutation of each gene does not affect viability and adults do not develop any overt phenotype. However, altered calcium flux was observed in *desma*-KO myofibres. These results show that loss-of-function of desmin genes by mutations in the first coding exon result in a mild phenotype with no visible muscle degeneration but altered calcium flux, in contrast to morpholino-mediated knockdown models and the *desma*^*sa5*^ allele.

## RESULTS

### Zebrafish desmin genes

In contrast to the single mammalian *DES* gene, zebrafish have two desmin orthologs, *desma* (*desmin a*) and *desmb (desmin b)*, which are paralog genes located on chromosome 9 and 6, respectively. Zebrafish *desma* and *desmb* paralogs share 81% and 83% identity with human *DES* gene. Although a single transcript is known to be encoded by human *DES* gene, in ENSEMBL two isoforms are predicted to be transcribed from zebrafish *desma*; *desma-1* mRNA coding for a 488 amino acid protein (predicted molecular weight of 55.7 kDa) and *desma-2* mRNA coding for a 473 amino acid protein (predicted molecular weight of 54.1 kDa). In *desma-2* transcript, exon 9 is skipped, corresponding to 15 amino acids located in the tail domain of Desma-1 protein (Supplementary Fig. S1 online). The 473 amino acid Desmb protein is predicted to have a molecular weight of 54.2 kDa. At the amino acid level, zebrafish Desma-1, Desma-2 and Desmb show 80%, 82% and 83% similarity with human desmin protein.

### Differential expression of desmin transcripts during zebrafish development

We analyzed the spatiotemporal expression patterns of *desma* and *desmb* genes during zebrafish development at several stages beginning from somitogenesis until 5 days post fertilization (dpf) by whole mount *in situ* hybridization (ISH). For detecting both *desma* transcripts (*desma-2* lacks exon 9), we synthesized an antisense probe by using a forward primer binding to exon 1 and a reverse primer binding to exon 10 (Supplementary Fig. S1 online). No specific staining was observed in embryos treated with sense probes for each gene (Supplementary Fig. S2 online). *Desma* was first detected at 11 hpf in adaxial slow muscle precursor cells when the first somites form and remained strongly expressed in somites at all examined stages (Fig. 1a). *Desma* was also expressed in the developing heart from 35 hpf (Fig. 1b). Cryosections of 72 hpf embryos showed that *desma* was distributed in the entire somite, including epaxial and hypaxial muscles (Fig. 1c). By 72 hpf, *desma* was expressed in cranial muscles including external ocular muscles, opercular muscles and mandibular muscles (Fig. 1d). At 96 hpf, transversal sections revealed *desma* expression at the anterior intestine (Fig. 1d, upper panel). At 72, 96 and 120 hpf, expression of *desma* in pectoral fin muscles was clearly visible (Fig. 1d). *Desmb* had an overlapping expression pattern to *desma* from the beginning of somitogenesis until 72 hpf, including staining in trunk somites and heart (Fig. 1a, b). At 72 hpf, somitic expression of *desmb* dramatically decreased with a residual expression in the lateral edges of the myotome. By contrast, strong signal of *desmb* transcripts was detected around the gut throughout the length of the intestine (Fig. 1a, c, e). Similar to *desma, desmb* was also expressed in pectoral fin muscles, operculum muscles and mandibular muscles at 72 hpf (Fig. 1e). These results indicate that *desma* and *desmb* expression partially overlap during zebrafish muscle development. As the gut develops, *desmb* expression shifts from somitic muscle to intestinal smooth muscle.

**Fig. 1.**
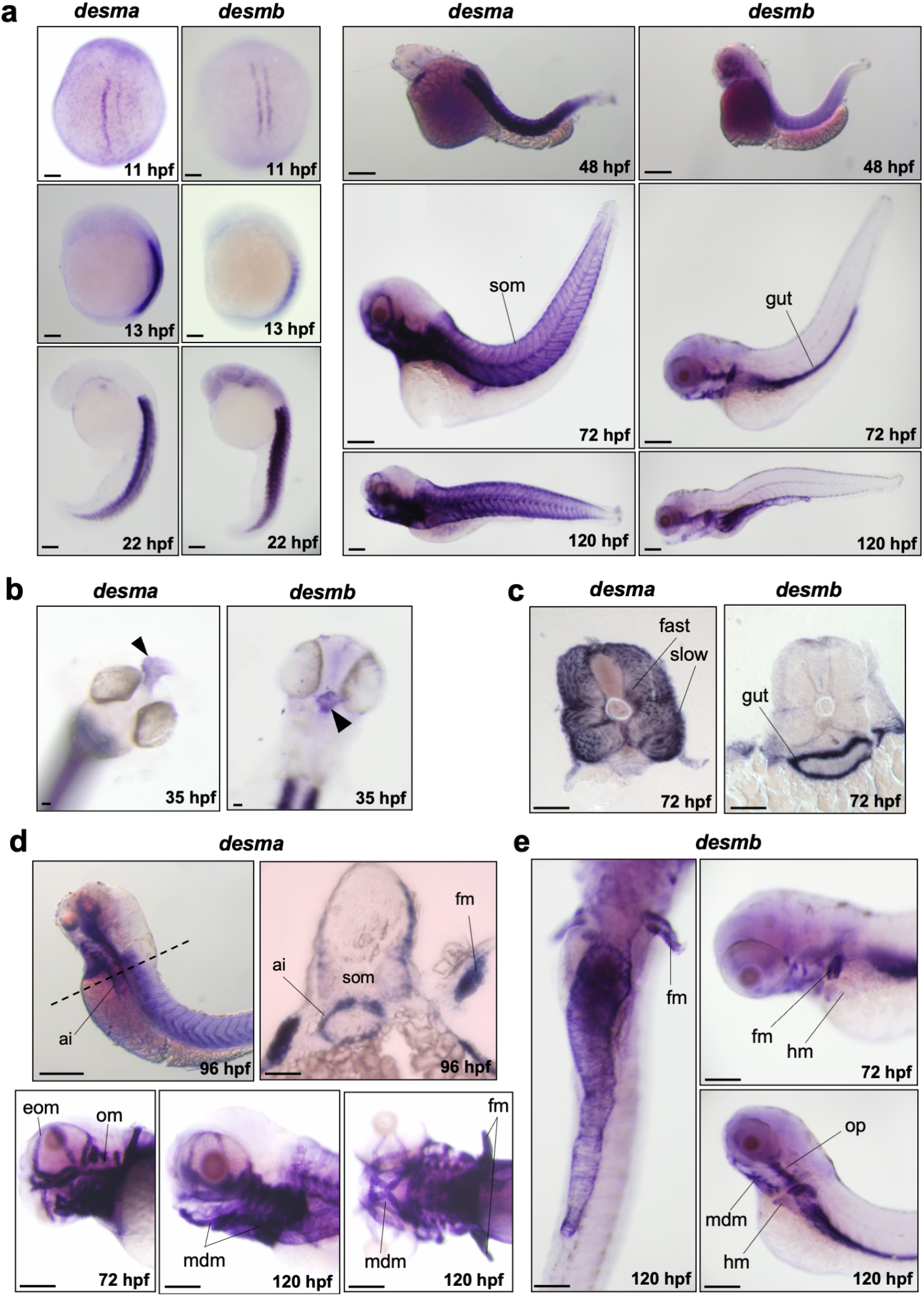
Whole mount *in situ* mRNA hybridisation of embryos at the indicated stages for antisense probes to *desma* and *desmb*. (**a**) Lateral views are anterior to top, dorsal to left for 13-22 hpf; anterior to left except 11 hpf which is a dorsal view. 48-120 hpf whole mounts are anterior to left, dorsal to top. Scale bar: 250 µm. (**b**) Arrows indicate heart in frontal view for left panel (*desma*) and ventral view for right panel (*desmb*) of 35 hpf embryos. (**c**) Transversal sections where dorsal is top. (**d**) Upper left panel is a lateral view at 96 hpf, scale bar: 250 µm. Dashed line represents the position of the transversal section in upper right panel where dorsal is top, scale bar: 100 µm. Lower panels are lateral views of zebrafish heads at 72 and 120 hpf treated with *desma* probe. Scale bar: 100 µm. (**e**) Left panel is a ventral view with anterior at top. Right panels are lateral views. Ai, anterior intestine; eom, extraocular muscles; fast, fast muscles; fm, pectoral fin muscles; hm, hypaxial muscles; mdm, mandibular muscles; om, opercular muscles; slow, slow muscles; som, somites. Scale bar: 250 µm.

### Generation of *desma* and *desmb* knockout zebrafish lines

*Desma* and *desmb* mutant zebrafish lines were generated by using CRISPR/Cas9 genome editing. The *desma*^*kg97*^ line exhibited 2 bp deletion and 4 bp insertion in the first exon (g.198_99delGAinsTGAT, NC_007120.7) leading to a frameshift at amino acid 45 and a premature stop codon after four amino acids. In the *desmb*^*kg156*^ mutant line, a 5 bp deletion in exon 1 (c.245_249delCTTAT, NM_001077452.1) resulted in a frameshift at amino acid 82 and introduced a premature stop codon after three amino acids (Fig. 2a). Heterozygous mutants showed no defect and homozygous mutant embryos and larvae developed relatively normally, at least for the first few days (Fig. 2d).

**Fig. 2.**
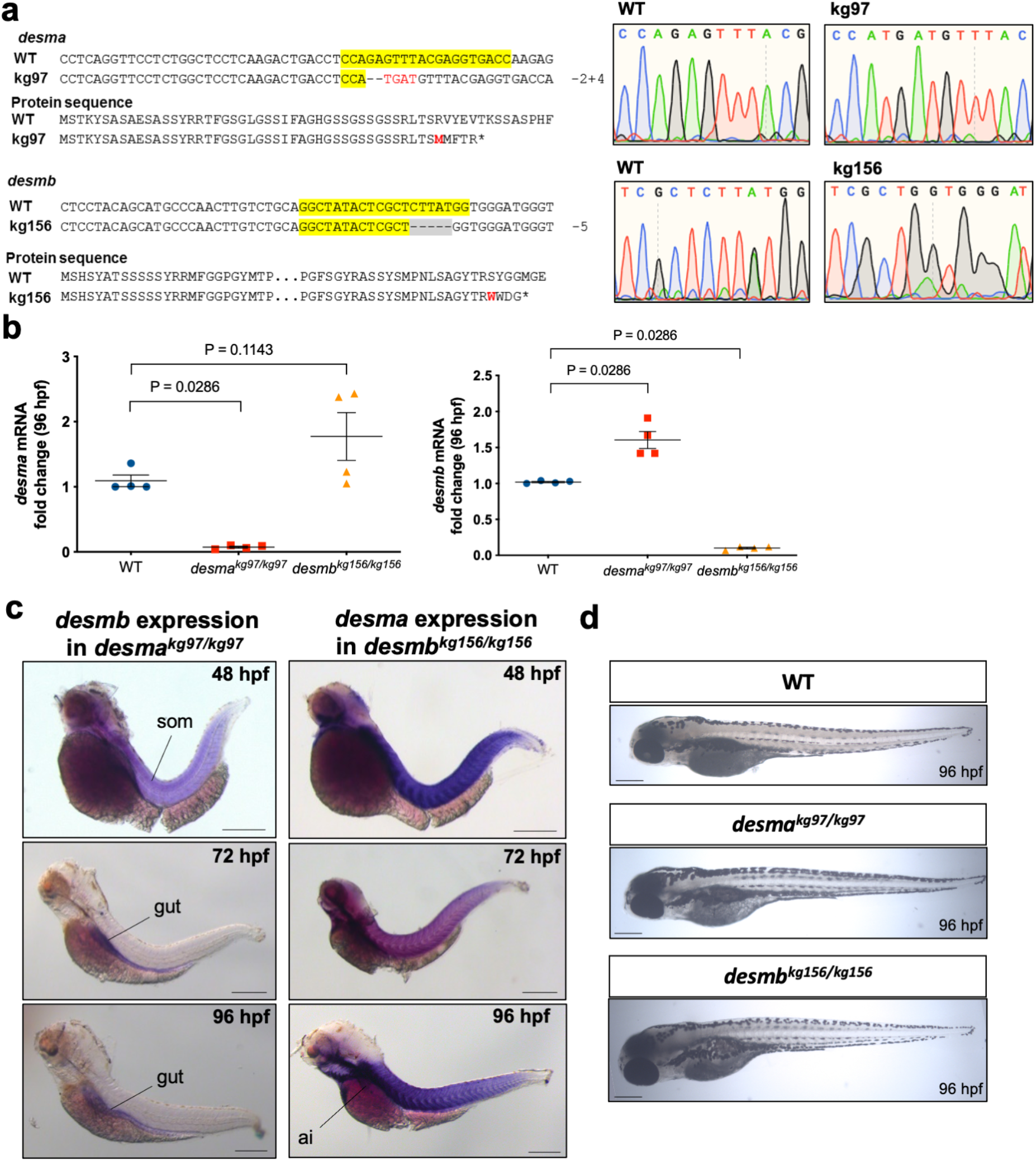
Generation of *desma* and *desmb* knockout lines. (**a**) Alignments and chromatograms of wild-type DNA sequences with mutant alleles, and predicted mutant polypeptide sequences. In DNA sequences, yellow highlights gRNA target sequence, hyphens show deleted bases, inserted bases are indicated in red font. In protein sequences, the first residue affected by the frameshift is indicated in red font, asterisks represent early stop codons. (**b**) Quantitative real-time PCR results showing the expression of *desma* and *desmb* mRNA in wild-type and homozygous mutant 96 hpf embryos (N=4, Mann-Whitney U). (**c**) Left panel shows whole mount *in situ* mRNA hybridisation of *desma*^*kg97*^ homozygous embryos at the indicated stages for antisense probes to *desmb*. Right panel shows whole mount *in situ* mRNA hybridisation of *desmb*^*kg156*^ homozygous embryos at the indicated stages for antisense probes to *desma*. Scale bar: 250 µm. Ai, anterior intestine; som, somite. (**d**) Brightfield pictures of 96 hpf WT and mutant embryos. Scale bar: 250 µm.

### Expression of *desma* and *desmb* in knockout lines

Mutations were predicted to trigger nonsense-mediated decay of mRNA or synthesis of short truncated polypeptides. As expected, qRT-PCR revealed a significant decrease of *desma* transcripts in *desma*^*kg97*^ homozygous embryos (*P*=0.0286, Mann-Whitney U), while *desmb* transcripts were significantly decreased in *desmb*^*kg156*^ homozygous embryos at 96 hpf compared to wild type (WT) (*P*=0.0286, Mann-Whitney U) (Fig. 2b). Consistent with El-Brolosy et al.^15^, weak *desmb* or *desma* mRNA up-regulation may occur in the respective mutant embryos (Fig. 2b).

In order to investigate the potential compensatory effect of *desmin* paralog upregulation in mutant embryos, expression patterns of *desmb* and *desma* transcripts were determined in *desma*^*kg97*^ or *desmb*^*kg156*^ homozygous mutant embryos, respectively. In *desma*^*kg97/kg97*^ embryos, the expression pattern of *desmb* transcripts (Fig. 2c) was similar to that of wild-type embryos (Fig. 1a). Somitic expression of *desmb* was observed in *desma*^*kg97/kg97*^ at 48 hpf, and a shift towards gut expression after 72 hpf. Beyond 72 hpf, no *desmb* staining was detected in somites of *desma*^*kg97/kg97*^ embryos. In *desmb*^*kg156/kg156*^ embryos, *desma* was still strongly expressed in somites and visible in the anterior intestine at 96 hpf (Fig. 2c). In order to confirm the loss-of-function of desmin in mutants at the protein level, expression of Desma and Desmb in WT, *desma*^*kg97*^ or *desmb*^*kg156*^ homozygous embryos at 96 hpf was determined by immunofluorescence staining using a desmin antibody recognizing both Desma and Desmb (Fig. 3). In WT embryos, desmin staining was detected in both somites and gut. As expected from ISH results, somitic expression was lost and gut was preferentially stained in *desma*^*kg97/kg97*^ mutants. In *desmb*^*kg156/kg156*^ mutants, expression in somites and anterior intestine were preserved while staining of the middle and posterior intestine was lost. We conclude that mutation of each gene reduces the cognate mRNA and protein at all stages examined and that compensation by up-regulation of the unmutated paralogous gene is at best very weak.

**Fig. 3.**
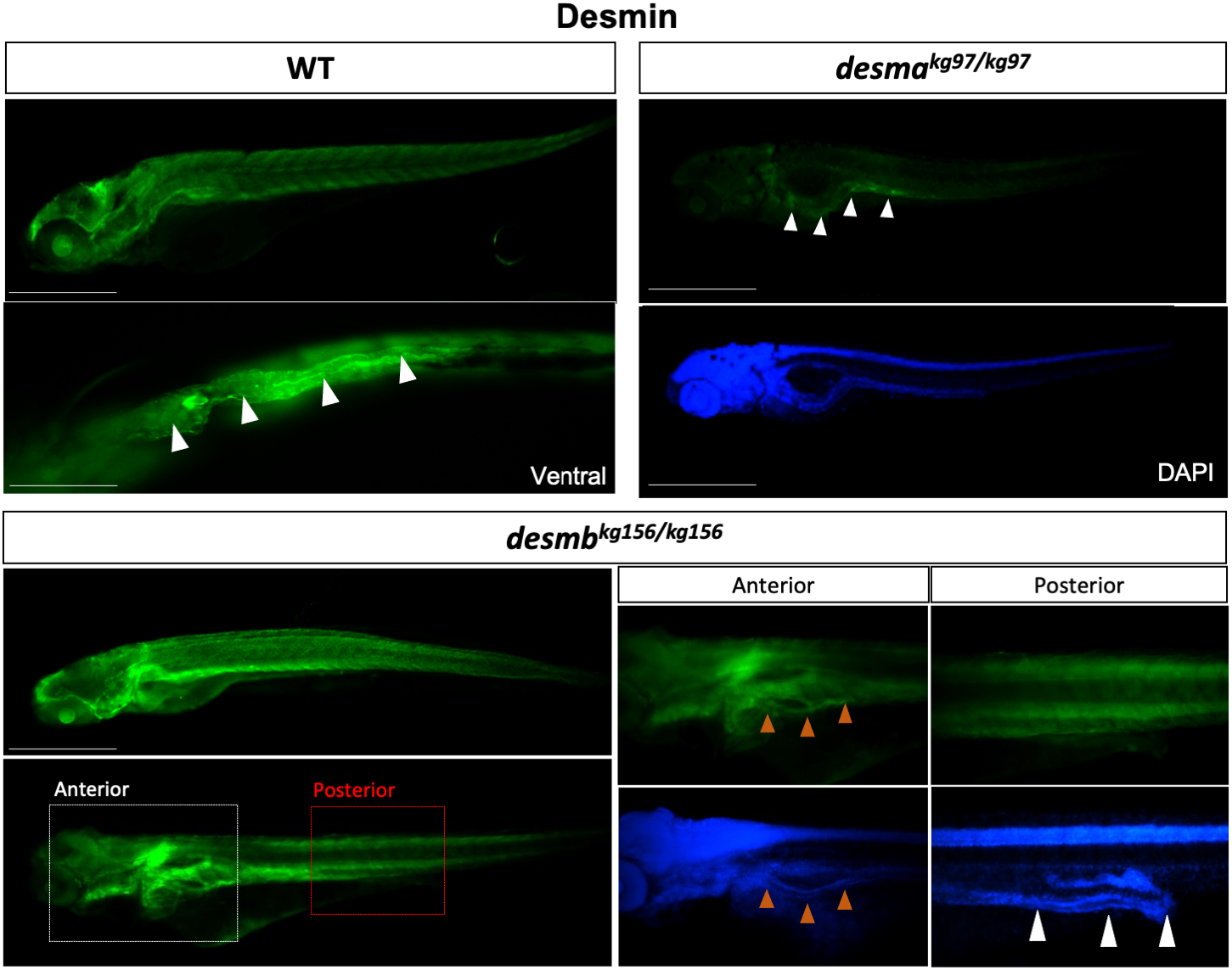
Desmin protein expression in 96 hpf embryos. Whole mount immunofluorescence staining of desmin with an anti-desmin polyclonal antibody (Sigma, D8281) recognizing both Desma and Desmb, in wild-type and mutant 96 hpf embryos. Left upper panel is lateral view of a WT embryo with anterior to left, dorsal to top, beneath is ventral view with anterior to left, white arrowheads indicate the gut. Right upper panels are lateral views of *desma*^*kg97/kg97*^ with anterior to left, dorsal to top. Lower panel shows *desmb*^*kg156/kg156*^ embryos with anterior to left, dorsal to top. Scale bar: 500 µm. Boxes with dashed lines delineate areas that were zoomed and focused on anterior and posterior segments of the intestine as shown in lower right panels, orange arrowheads indicate anterior intestine, white arrowheads show middle and posterior segments of the intestine.

### General larval characteristics of desmin mutants

In order to evaluate the effects of the absence of *desma* or *desmb* on development, WT, *desma*^*kg97*^ and *desmb*^*kg156*^ homozygous embryos/larvae were compared. No significant difference was observed in the number of viable eggs between *desma*^*kg97/kg97*^ and WT (*P*=0.6842, Mann-Whitney U) or *desmb*^*kg156/kg156*^ and WT (*P*=0.9654, Mann-Whitney U) (Fig. 4a). Hatching period (48-72 hpf^16^) is a critical process in embryonic development and reduction in the hatching rate could be associated with reduced muscle function^17^. The time course showed that although different hatching times were observed between groups, no significant difference was found between hatching rates of mutant and WT embryos at 72 hpf with over 94% of embryos hatched (*desma*^*kg97/kg97*^ vs. WT, *P*=0.9902; *desmb*^*kg156/kg156*^ vs WT, *P*=0.3186, repeated measures two-way ANOVA, Bonferroni post *hoc* test) (Fig. 4b).

**Fig. 4.**
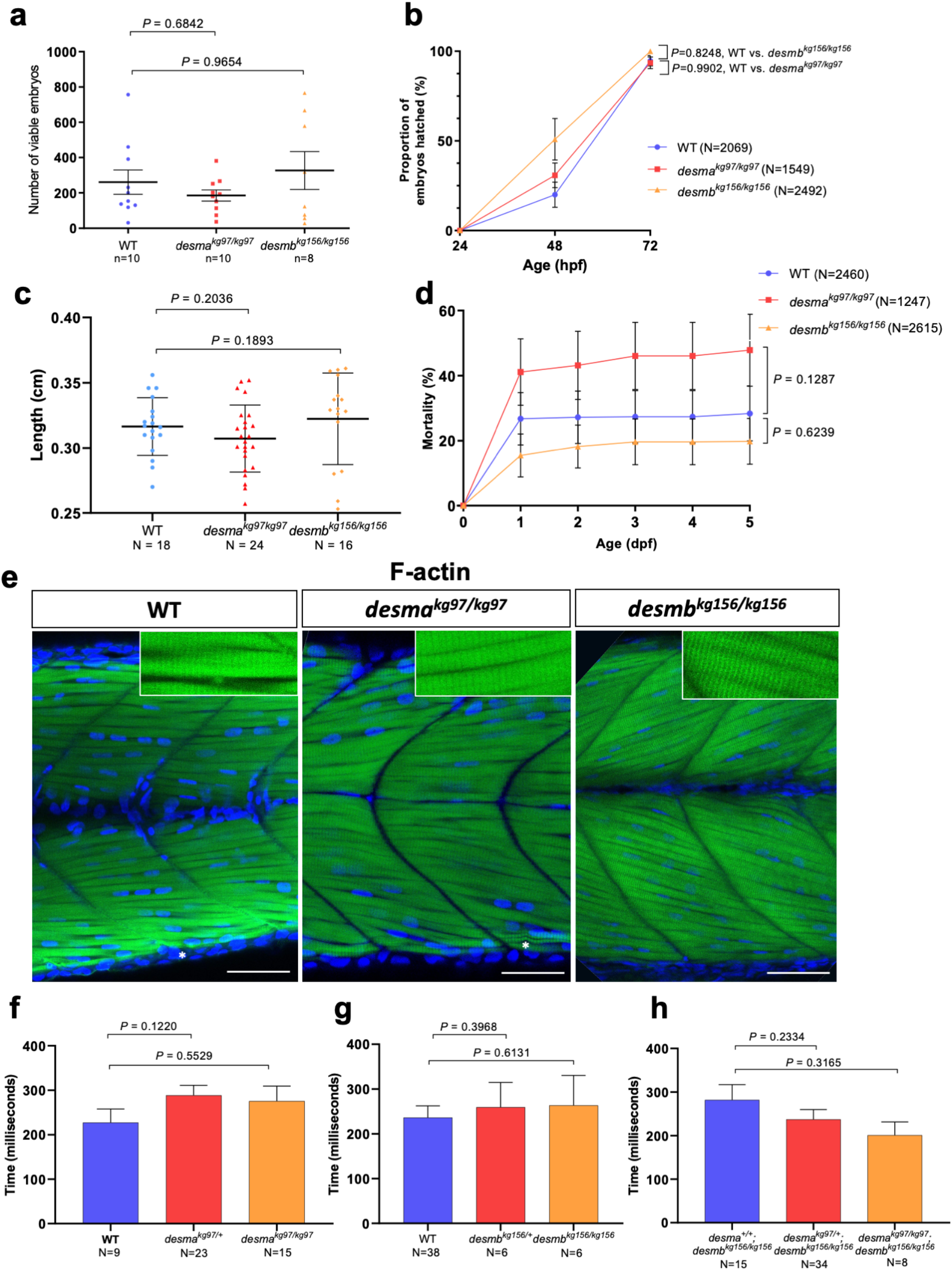
Evaluation of neuromuscular defect in mutant larvae. (**a)** The number of viable embryos after successful mating of WT fish (n=10) compared to homozygous *desma*^kg97^ (n=10) (*P*=0.6842, Mann-Whitney U) or homozygous *desmb*^kg156^ (n=8) (*P*=0.9654, Mann-Whitney U) fish. (**b)** Comparison of hatching rate between homozygous mutants and WT 24, 48 and 72 hpf embryos At 72 hpf, no significant difference in the hatching rate was found between mutants and WT (*desma*^*kg97/kg97*^ vs. WT, *P*=0.9902; *desmb*^*kg156/kg156*^ vs WT, *P*=0.3186, repeated measures two-way ANOVA, Bonferroni post *hoc* test) (**c)** Body length of 96 hpf WT (N=18) and homozygous *desma*^*kg97*^ (N=25) or homozygous *desmb*^*kg156*^ (N=16) mutant embryos (*P*=0.2036 for WT vs. *desma*^*kg97/kg97*^; *P*=0.1893 for WT vs. *desmb*^*kg156/kg156*^, Mann-Whitney U). (**d)** Cumulative mortality rate from 1 to 5 dpf homozygous *desma*^*kg97*^ mutants (N=1247) compared to WT (N=2460) (*P*=0.1287, repeated measures two-way ANOVA, Bonferroni post *hoc* test) and homozygous *desmb*^*kg156*^ mutants (N=2615) compared to WT (*P*=0.6239, repeated measures two-way ANOVA, Bonferroni post *hoc* test). **(e)** Optical sections of the mid-trunk region of WT and homozygous mutants 96 hpf embryos (N=8) stained with rhodamine phalloidin (F-actin) with insets showing striations (asterisks indicate occasional wavy fibres in both WT and mutants). Scale bar: 50 µm. (**f-h)** Touch-evoked escape time (ms) of 48 hpf embryos. Motility experiments were performed blind on siblings from (**f**) *desma*^*kg97/+*^, (**g**) *desmb*^*kg156/+*^ or (**h**) *desma*^*kg97/+*^*;desmb*^*kg156/kg156*^ in-crosses followed by *post hoc* genotyping. (**f**) Heterozygous *desma*^*kg97*^ embryos (N=23) were compared to WT (N=9) (P=0.122, Mann-Whitney U) and *desma*^*kg97/kg97*^ (N=15) compared to WT (P=0.5529, Mann-Whitney U). (**g**) Heterozygous *desmb*^*kg156*^ embryos (N=6) were compared to WT (N=38) (P=0.3968, Mann-Whitney U) and *desmb*^*kg156/kg156*^ (N=6) compared to WT (P=0.6131, Mann-Whitney U). (**h**) Homozygous double mutants (N=8) were compared to *desma*^+/+^;*desmb*^*kg156/kg156*^ (n=15) (*P*=0.3165, Mann-Whitney U) and *desma*^*kg97/+*^*;desmb*^*kg156/kg156*^ embryos (n=34) compared to *desma*^+/+^;*desmb*^*kg156/kg156*^ (*P*=0.2334, Mann-Whitney U).

Similar mortality rates were observed with no significant difference between homozygous mutants of *desma*^*kg97*^ or *desmb*^*kg156*^ and WT embryos (*desma*^*kg97/kg97*^ vs. WT, *P*=0.1287; *desmb*^*kg156/kg156*^ vs. WT, *P*=0.6239, repeated measures two-way ANOVA) (Fig. 4d). Finally, among surviving larvae, no statistically significant difference in body length was observed between WT and mutant groups (*P*=0.2036 for WT vs. *desma*^*kg97/kg97*^; *P*=0.1893 for WT vs. *desmb*^*kg156/kg156*^, Mann-Whitney U) (Fig. 4c).

### Mutant larvae show no neuromuscular defect

Muscle fibre integrity and somite morphology length were investigated by staining muscle actin with rhodamine phalloidin in 96 hpf embryos (Fig. 4e). At this stage, mainly *desma* is expressed in somites. No muscle lesion or detachment of fibres was observed in mutants and no significant difference was found in somite length, fibre diameter and nuclei number between mutants and WT (Supplementary Fig. S3 online).

Touch-evoked response assay is a widely used method to assess neuromuscular function in 48 hpf zebrafish embryos^18^. We performed motility experiments in a blinded manner on siblings from three groups of heterozygous mutant in-crosses and genotyped them after. First, the average escape time of siblings from *desma*^*kg97/+*^ in-crosses was assessed and no significant difference was found in *desma*^*kg97*^ heterozygous (*P*=0.122, Mann-Whitney U) or homozygous (*P*=0.5529, Mann-Whitney U) mutants compared to WT (Fig. 4f). Similarly, no significant difference in the average escape time was found in *desmb*^*kg156*^ heterozygous (*P*=0.3968, Mann-Whitney U) or homozygous (*P*=0.6131, Mann-Whitney U) mutants compared to WT (Fig. 4g). Since both *desma* and *desmb* are expressed in somites up to this stage (48 hpf) and might potentially compensate for the absence of each other, we finally evaluated the effect of the absence of both proteins in siblings from *desma*^*kg97/+*^*;desmb*^*kg156/kg156*^ double mutants. No significant difference in the average escape time was found in homozygous double mutants (*desma*^*kg97/kg97*^*;desmb*^*kg156/kg156*^) compared to *desma*^*+/+*^*;desmb*^*kg156/kg156*^ (*P*=0.3165, Mann-Whitney U) and in *desma*^*kg97/+*^*;desmb*^*kg156/kg156*^ embryos compared to *desma*^*+/+*^*;desmb*^*kg156/kg156*^ (*P* = 0.2334, Mann-Whitney U) (Fig. 4h).

### Mutant adult skeletal muscle shows no muscle degeneration

Because no skeletal muscle phenotype was observed in mutant larvae, surviving one-year-old adults were investigated for muscle defects. Some homozygous single and double mutants survived until adulthood with no sign of behavioral defect (Fig. 5a). Western blot was performed using myotomal skeletal muscle protein extracts isolated from adult fish (WT, homozygous and heterozygous *desma*^*kg97*^ mutants, homozygous and heterozygous *desmb*^*kg156*^ mutants and *desma*^*kg97/kg97*^*;desmb*^*kg156/kg156*^ double mutants). As *desma* was shown to be predominantly expressed in somites from 72 to 120 hpf, the two bands detected in WT and *desmb*^*kg156*^ mutants were predicted to correspond to Desma-1 (predicted molecular weight 55.7 kDa) and Desma-2 (predicted molecular weight 54.1 kDa) isoforms. In agreement with that, both bands were absent in *desma*^*kg97*^ homozygous mutant and double mutants while still expressed in homozygous *desmb*^*kg156*^ mutant (Fig. 5b). No *desmb* mRNA upregulation was observed in *desma*^*kg97/kg97*^ adult skeletal muscle, nor significant *desma* mRNA upregulation in *desmb*^*kg156/kg156*^ compared to WT. Moreover, *desmb* mRNA expression was >1200-fold lower than *desma* in WT adult skeletal muscle tissue, confirming that Desma is the predominantly expressed paralog in adult skeletal muscle tissue (Supplementary Fig. S4 online).

**Fig. 5.**
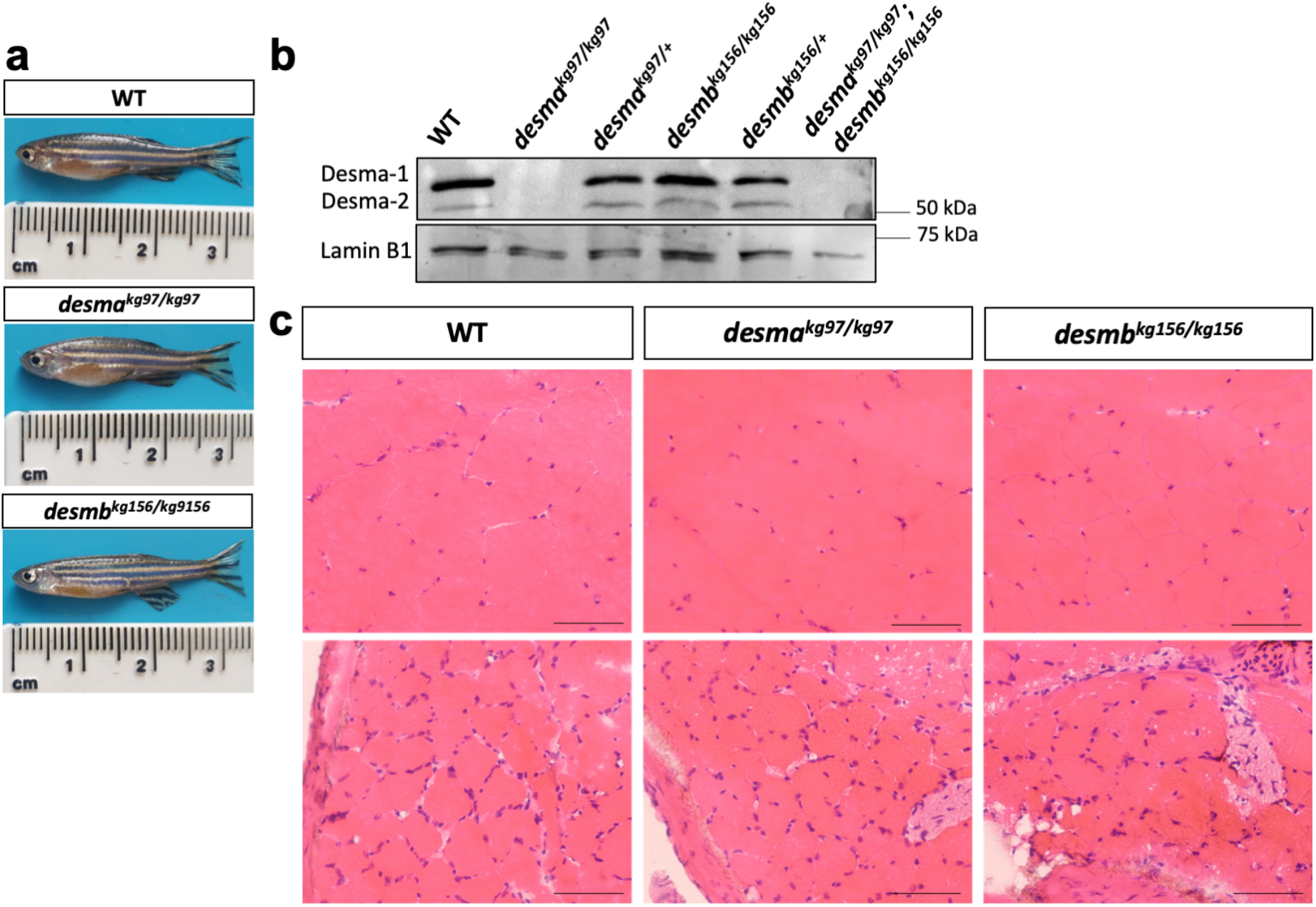
Desmin protein expression and histological examination of adult skeletal muscle. (**a)** Photographs of 1 yr-old anesthetized fish. (**b**) Immunoblotting of skeletal muscle protein extracts of WT, *desma*^*kg97/kg97*^, *desma*^*kg97/+*^, *desmb*^*kg156/kg156*^, *desmb*^*kg156/+*^ and *desma*^*kg97/kg97*^*;desmb*^*kg156/kg156*^ double mutant fish probed with anti-desmin antibody. Lamin B1 was labeled on the same blot as loading control. Uncropped full-length pictures of the blot is available in Supplementary Figure S5 online. (**c)** Transversal cryosections of mid-trunk somites from two different areas of WT and homozygous mutant adults stained with hematoxylin eosin showing no pathological changes. Scale bar: 25 µm.

Histopathological examination of hematoxylin-eosin stained skeletal muscle tissue sections revealed no pathological changes in homozygous single mutants compared to WT (Fig. 5c). General morphology of skeletal muscle fibre shape and size, integrity of sarcoplasm and interstitium were normal. Mild variation in fibre size was present in both WT and mutant fish. Signs of degeneration or regeneration, internal nuclei, inflammatory cell infiltration and increase in connective or adipose tissue in interstitium were absent (Fig. 5c).

Transmission electron microscopy revealed intact sarcolemma, generally peripherally, unfrequently centrally located healthy euchromatic nuclei with well determined nucleoli and perinuclear cisternae in adult WT, *desma*^*kg97/kg97*^ and *desmb*^*kg156/kg156*^ skeletal muscle tissue samples (Fig. 6a-f). WT and both mutant samples presented pleomorphic mitochondria with well-defined tubular cristae. Mitochondria making rows or clusters and exhibiting fission and mitophagy were common in all groups (Fig. 6g-l). The sarcoplasmic reticulum and T-tubule system-associated triads appeared locally proliferated and enlarged in all samples (Fig. 6p-r). These membrane systems caused the shift of the sarcomeric organization to some extent in both WT and mutant samples (Fig. 6a-c, m-r). Ultrastructural evaluation revealed a similar myofilament organization in all group’s muscle samples. The Z lines were generally aligned in most of the fibres in all groups (Fig. 6a-c). A limited number of Z line streaming or duplication was noted in both WT and knockout muscle specimens (Fig. 6m-o). As expected, no deposit or particle was noted in any of the groups. We conclude that fish entirely lacking wild-type Desma generate and maintain functional skeletal muscle in the context of a zebrafish aquarium.

**Fig. 6.**
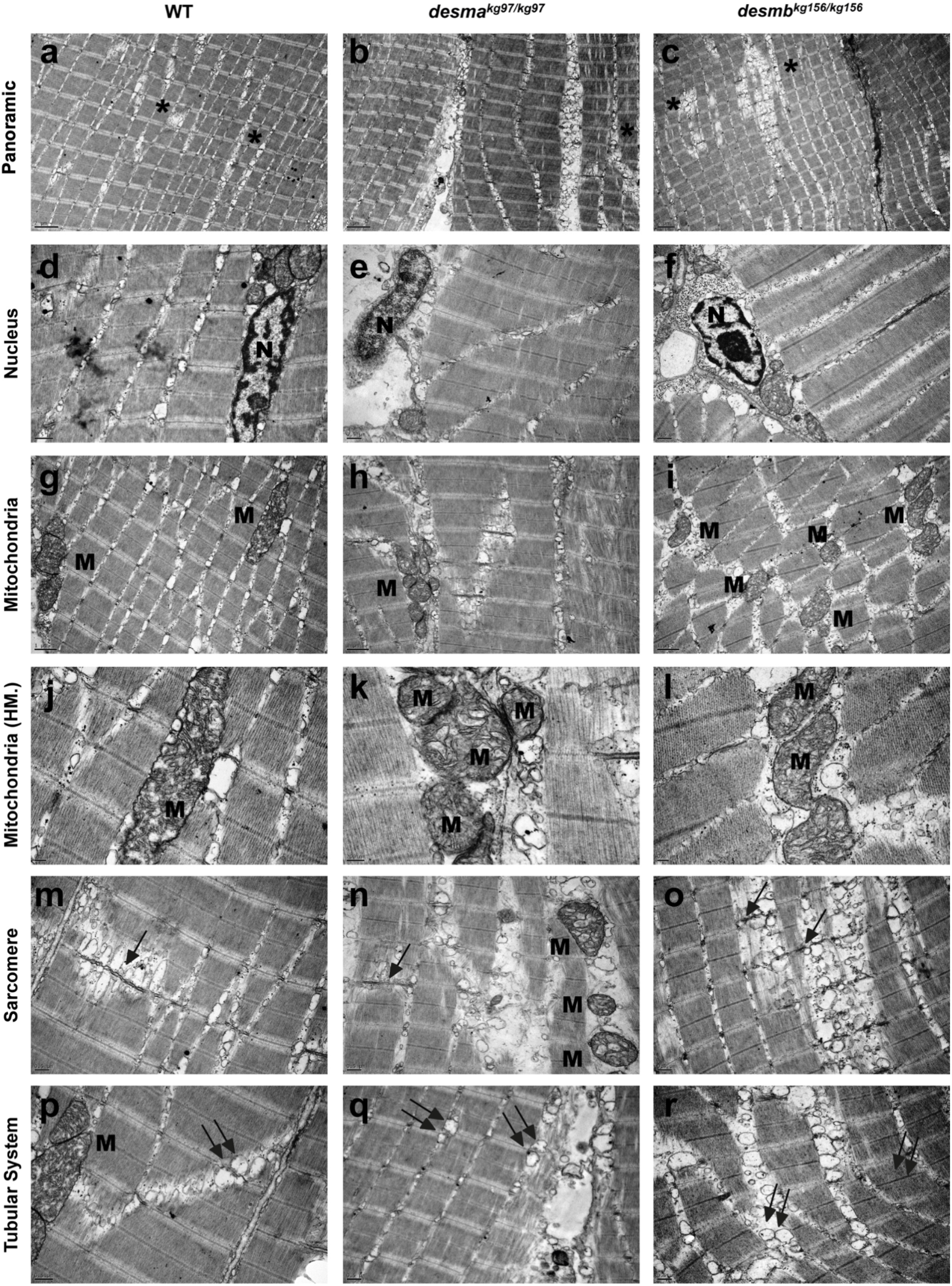
Ultrastructural features of adult WT and mutant skeletal muscle tissue. First, second and third columns present adult WT, *desma*^*kg97/kg97*^, *desmb*^*kg156/kg156*^ skeletal muscle tissues’ electron micrographs, respectively. First row indicates the generally aligned sarcomeric structures with the presence of seldom disruptions (asterisks). Second row shows centrally (d) and peripherally (e, f) located healthy nuclei (N) with peripherally condensed chromatin and well-defined nucleoli. Third and fourth row (g-l) shows pleomorphic mitochondria (M) with well-defined tubular cristae. Fifth and sixth rows present the Z line streaming (arrows in m-o) and enlargement (m-r) of the T-tubule systems related to the triads (double arrows). Uranyl acetate, lead citrate. HM, High Magnification. Magnifications are as follows: (a-c) 8000x, (d) 25000x, (e,f) 20000x, (g-i) 15000x, (j) 40000x, (k) 60000x, (l) 30000x, (m-o) 20000x, (p) 25000x and (q, r) 20000x.

### Altered calcium flux in *desma* mutant fibres

Although anatomically apparently wild-type, adult *desma*^*kg97/kg97*^ mutants were physiologically defective in calcium handling. To investigate amplitude and time course of calcium signals released in response to a depolarizing voltage stimulus, individual fibres were dissected from WT (N=8), *desma*^*kg97*^ (N=11) or *desmb*^*kg156*^ (N=8) homozygous mutant 1-year-old fish. Calcium flux along fibres was monitored in the isolated fibres by Fluo-4 AM after four consecutive depolarizing stimuli with the amplitude of the current pulse kept constant at 100 nA and the duration increased by 10 ms at each pulse (Fig. 7a-b). Amplitudes of the calcium emission signals in the first (10 ms) stimulus and fourth (40 ms) stimulus were compared between mutants and WT. To eliminate variance induced by amplitude, offset and fibre size differences, amplitude values were baseline corrected and divided by the diameter of each fibre. Amplitudes of the first and the fourth calcium transient were significantly lower in *desma*^*kg97/kg97*^ fibres compared to WT fibres (*P*=0.0008 for first transient; *P*=0.0025 for fourth transient, Mann-Whitney U) (Fig. 7c-d). In contrast, no significant difference was found in the amplitude of calcium flux between *desmb*^*kg156/kg156*^ fibres and WT fibres (*P*=0.0881 for first transient; *P*=0.6657 for fourth transient, Mann-Whitney U) (Fig. 7c-d).

**Fig. 7.**
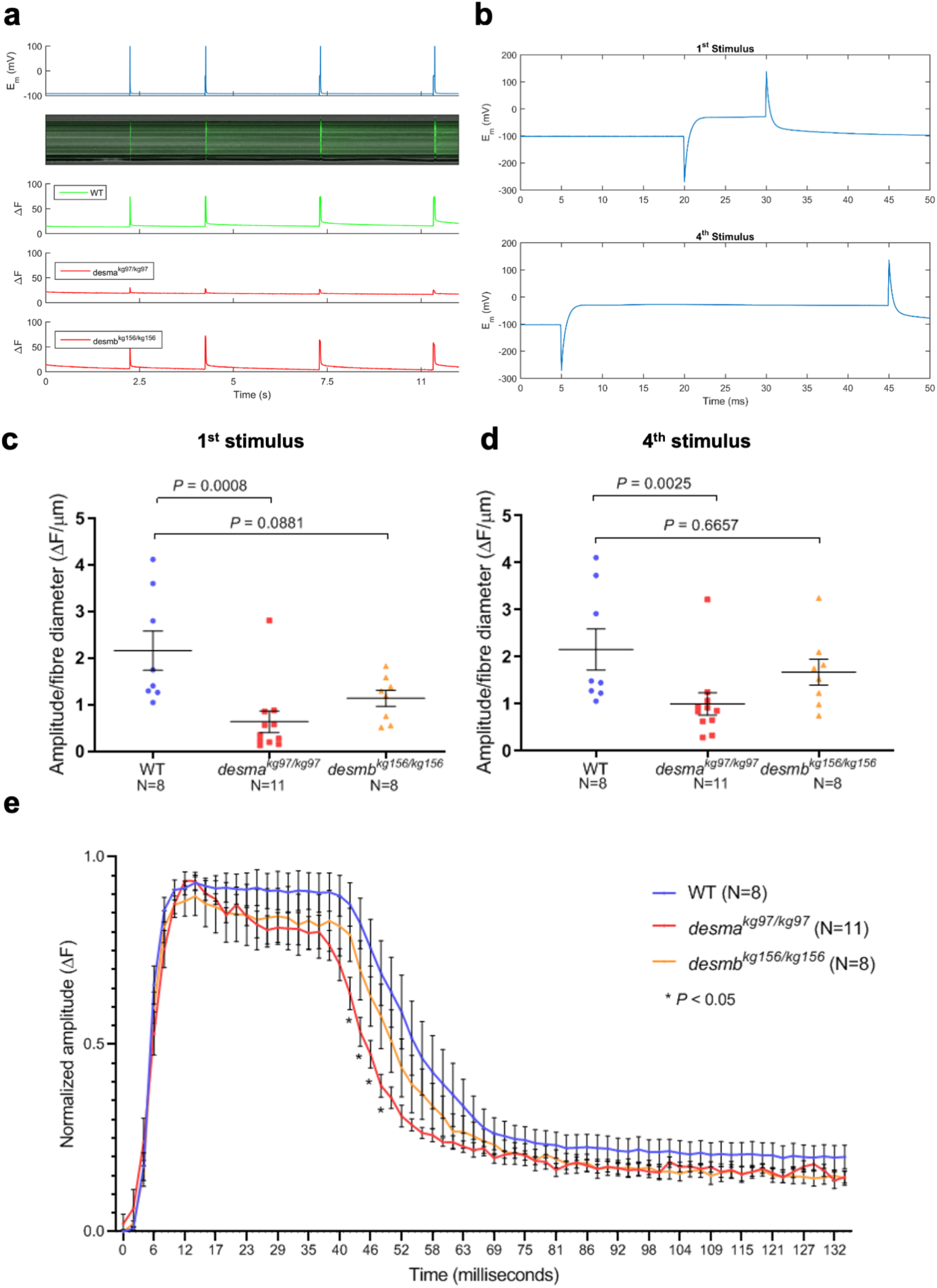
Calcium flux in isolated fibres. (**a**) Calcium flux along fibres was monitored in vivo by Fluo-4 AM after four consecutive depolarizing stimuli. Top panel, membrane potential responses to current stimulus pulses with varying durations of 10, 20, 30, 40 ms with the amplitude of the current pulse kept constant at 100 mV and the duration increased by 10 ms at each pulse. Middle panel, evoked calcium transients in line scan mode (*tx*) to current stimuli. Bottom panel, integrated emission signal as a function of time for each experimental group. (**b**) Membrane potential response to the first stimulus of 10 ms (top) or to the fourth stimulus of 40 ms. (**c**) Baseline corrected amplitude divided by fibre diameter (ΔF/µm) of the calcium emission signals during the first (10 ms) stimulus and fourth (40 ms) stimulus were compared between mutants and WT. (**e**) Time course analysis of the baseline corrected and normalized amplitude of the calcium transient from the longest stimulus (40 ms) was represented as mean amplitude values as a function of time. Homozygous *desma*^*kg97*^ fibres (N=11, red curve) or homozygous *desmb*^*kg156*^ fibres (N=8, orange curve) were compared to WT (N=8, blue curve). (* indicates time points where *P*<0.05, repeated measures two-way ANOVA, Bonferroni post *hoc* test).

Time course of the longest calcium transient (40 ms, fourth transient) has been analyzed for comparing the waveform of the responses and dissecting the phases of the transient. Signals have been baseline corrected and then normalized. Mean amplitude values as a function of time were compared between *desma*^*kg97/kg97*^ and WT or *desmb*^*kg156/kg156*^ and WT. No significant difference was found in the rising and the plateau phases of the recorded transients between the experimental groups (Figure 7e). However, the decay phase was significantly faster in *desma*^*kg97/kg97*^ mutant fibres compared to WT (* indicates *P*<0.05, repeated measures ANOVA, Bonferroni post *hoc* test). No significant difference was found in *desmb*^*kg156/kg156*^ mutant fibres compared to WT (Fig. 7e).

## DISCUSSION

Although previous studies showed that *desma* is the main gene expressed in somites, no study did clearly distinguish spatial and temporal differences in the expression of the two genes. We showed that until 72 hpf *desma* and *desmb* expressions overlap in somites and heart but later in development, *desmb* expression shifts from somites to intestinal smooth muscle, and *desma* remains as the predominantly expressed paralog in adult somites. Note that the residual *desmb* expression detected in the lateral edges of the myotome might reflect a transient expression in new skeletal muscle fibres (Fig. 1c)^19^. *In silico* prediction of transcription factor (TF) binding sites on *desma* and *desmb* putative promoters revealed 14 and 7 muscle-related TFs unique to *desma* and *desmb*, respectively (Supplementary Table S2 online). TFs exclusively binding to *desma* promoter were enriched in striated muscle (12 out of 14 TFs) while TFs unique to *desmb* promoter were enriched in smooth muscle (5 out of 7 TFs). This preliminary data supports the hypothesis that *desma* and *desmb* expressions are differentially regulated throughout zebrafish muscle development and experimental confirmation would bring further insight into their transcriptional regulation. Lack of desmin in mice affects multiple tissues including myocardium, skeletal muscle, intestinal smooth muscle and smooth muscle of the aorta^8^. Since Desmb appears to be the major Desmin in smooth muscle in zebrafish, at least in the gut, zebrafish *desmb* mutants may be particularly useful for understanding the role of desmin in smooth muscle without interference by additional phenotypes from skeletal muscle. It should be noted that vascular defects have been observed in *desma*^*sa5*^ mutant hearts where *desmb* expression is not affected; therefore, it would be interesting to investigate the expression of *desma* in early smooth muscle development and endothelial cells. Also, comparison of the filament-forming capacity of Desma and Desmb would offer further insight into their functional redundancy. Although not the focus of this study, it should be noted that heart and gastrointestinal development in mutant lines have not been extensively analyzed; however; no heart oedema such as observed in desmin morphant or overt morphological abnormality in the gut have been observed in *desma*^*kg97*^ or *desmb*^*kg156*^ embryos and adults.

Reduced *desma* or *desmb* mRNAs in their respective mutants suggest the mRNAs are degraded by nonsense-mediated decay, such that only tiny amounts of the severely truncated proteins would be produced. Such fragments lack the highly conserved α-helical central rod domain responsible for filament formation and all known protein-protein interaction domains of Desmin^20^. Intriguingly, no overt skeletal muscle phenotype was observed in desmin knockout mutants. These results are in contrast to those reported for desmin morphants where both *desma* and *desmb* were targeted with morpholinos^13^ and for ENU-induced *desma*^*sa*^ knockout mutants^21^. Both morphants and mutants embryos exhibited disorganized fibres, disrupted sarcomeres and altered motility. Our findings raise the possibility that a haploinsufficient dominant effect in *desma*^*sa5*^ mutants may disrupt muscle; such an explanation would no account for the morphant phenotype. We conclude that at least two further alleles of *desma* may be needed to resolve these differences: firstly, a full locus or promoter deletion allele that would simultaneously prevent any Desmin polypeptide synthesis and any RNA-triggered compensation and, secondly, an allele lacking morpholino binding ability but nevertheless encoding wild-type protein to prove that the morphant phenotype is due to *desma* binding.

Many loss-of-function mutants not exhibiting an overt phenotype and failing to phenocopy morphants have been previously reported^22–24^. El-Brolosy et al. proposed transcriptional adaptation triggered by mutant RNA degradation as a mechanism to explain this discrepancy^15,25^. According to this model, decay of the mutant mRNA results in upregulation of related genes, alleviating or suppressing the phenotype observed in morphants. Although a slight increase in *desmb* mRNA expression was observed in *desma*^*kg97*^ embryos, no *desmb* mRNA upregulation and Desmin protein expression have been detected in *desma*^*kg97*^ adult skeletal muscle, suggesting that the compensating gene in fish carrying our *desma* allele is not *desmb*, the only gene known to have high and extensive homology to *desma*. However, we found that vimentin mRNA level was upregulated by approximately 10-fold in *desmb*^*kg156*^ adult skeletal muscles (Supplementary Fig. S4 online). Vimentin and desmin are closely related members of the class III intermediate filament (IF) proteins, vimentin being the major IF protein expressed early in development and in mesenchymal cells while desmin remains as the main IF protein expressed in muscle^26^. In zebrafish embryos, vimentin expression is restricted to the developing brain and in adult tissues, vimentin mRNA and protein are only detected in skin and eye^27^. The mechanisms underlying vimentin upregulation in *desmb*^*kg156*^ mutants need further investigation. Genetic adaptation by yet unknown adapting genes could explain the milder phenotype observed in mutants created in our study compared to desmin morphants. However, it has been shown that *desma*^*sa5*^ allele results in a muscle phenotype similar to that of morphants, even though the *desma*^*sa5*^ allele predicts an in-frame stop codon and no *desma* mRNA was detected in mutants^14,21^. The specific mechanism leading to the absence of mRNA was not investigated either in *desma*^*sa5*^ mutants or in the present study. It is possible that the *desma*^*sa5*^ allele is not as efficient as *desma*^*kg97*^ allele in triggering an adaptation mechanism for yet unknown reasons. It will be interesting to compare these models in the contexts of mRNA decay and transcriptional adaptation. Alternatively, short truncated polypeptides not detected in Western blot might play a role in this process before being rapidly degraded. In conclusion, unraveling modifier genes and proteins that compensate for the loss of desmin in our models will bring further insights in the role of desmin in muscle.

We found that the amplitude of calcium signals released in response to a depolarizing current stimulus was significantly lower accompanied by a faster decay in *desma*^*kg97*^ skeletal muscle fibres compared to WT. In desmin-null mice, it was shown that the lack of desmin significantly altered the decay rate of the Ca^2+^ transient and depressed the speed of Ca^2+^ uptake into the sarcoplasmic reticulum which caused a rise in cytosolic Ca^2+^ concentration at rest^28^. Similarly, knockdown of desmin in rat cardiomyocytes decreased the Ca^2+^ concentration in the sarcoplasmic reticulum and has been related to inhibition of SERCA2 expression^29^. Impaired mitochondrial morphology, positioning and respiratory function have been reported in human and mice skeletal and cardiac muscles harboring various desmin mutations. Disruption of the inner mitochondrial membrane and cristae organization has been observed in desmin-deficient mice cardiomyocytes^30^. The contribution of mitochondrial calcium uptake in regulating calcium homeostasis in skeletal muscle cells remains poorly understood; however, several studies have linked the dysregulation of mitochondrial calcium uptake to altered skeletal muscle function^31,32^. Particularly, it was shown that aggregate-prone mutations in mice desmin gene cause a reduction in the capacity of mitochondrial calcium uptake upon both electrical and chemical stimulation of myotubes^33^. In zebrafish, disruption of mitochondrial cristae organization has been observed in skeletal muscle of an aggregation prone *desma*-knockin line while *desma*^*sa5*^ knockout mutants presented with normal mitochondria; however, defects in heart calcium flux were observed in both mutants and linked to mislocalization of ryanodine receptors^14^. Similarly, in our study, mitochondrial shape and structure was not affected in *desma*^*kg97*^ adult skeletal muscle tissue while calcium flux was altered. Further investigation is needed to gain insight into the mechanism underlying defective calcium signaling in desmin-null zebrafish mutants.

## METHODS

### Ethical approval and zebrafish maintenance

All animal procedures were performed in accordance with Turkish *Animal Protection Act* 2004 and later modifications conforming to all relevant guidelines and regulations, and approved by the Hacettepe University Animal Experimentations Local Ethics Board (Approval no. 2014/07-08). The work in the UK was performed under UK Home Office license in accordance with the UK *Animal (Scientific Procedures) Act 1986* and subsequent modifications. Adult zebrafish (AB) (*Danio rerio*) were kept in 14/10 hr light-dark cycle at 28.5°C. Adults are fed twice a day, with dry feed in morning and *Artemia spp*. in evening. For spawning, male and female fish were placed in a breeding tank with a separator. Next morning the separator was removed, viable eggs were collected and rinsed in E3 medium (5 mM NaCl, 0.17 mM KCl, 0.33 mM CaCl_2_, 0.33 mM MgSO_4_, 10^−5^% Methylene Blue). Embryos were kept in E3 medium for five days and placed into adult tanks containing system water. Embryos were fed five times a day with dry feed and Artemia.

### Whole mount *in situ* hybridization

Embryos were fixed in 4% PFA in PBS (w/v) overnight at 4°C, dehydrated in 50% and %100 methanol washes and stored at -20°C at least overnight before use. Probe templates were amplified from cDNA isolated from 96 hpf wild-type embryos. Reverse primers for antisense probe contained T3 promoter sequence while forward primers for sense probes contained T7 promoter sequence. Primers sequences can be found as Supplementary Table S1 online. Digoxigenin-labelled probes were generated using T3 or T7 RNA polymerases. Whole mount *in situ* mRNA hybridization was performed as described^34^. Embryos were photographed as on Zeiss Axiophot with Axiocam (Carl Zeiss, Oberkochen, Germany) using Openlab software (Agilent, Santa Clara, CA, USA).

### Generation of mutant zebrafish lines

CRISPR-Cas9 genome editing was performed as described in Fin et al^24^. CRISPR-Cas9 sgRNAs showing minimal off-target sites were designed by using the online software ZiFiT Targeter. Oligos encoding sgRNAs targeting the first exon of *desma* (GRCz11, Chr9: 7539113-7539132, GGTCACCTCGTAAACTCTGG) and *desmb* (GRCz11, Chr6: 13891417-13891437, GGCTATACTCGCTCTTATGG) were cloned into pDR274 plasmid (Addgene). sgRNAs were synthesized using T7 RiboMAX Large Scale RNA production system (Promega) following the manufacturer’s protocol. Transcribed sgRNAs were purified using sodium acetate and ethanol precipitation and quantified on Qubit fluorometer. Cas9 mRNA was transcribed from pCS2-Cas9 plasmid using mMessage mMachine SP6 Kit (Ambion) and purified by lithium chloride precipitation. One-cell stage embryos were injected with 2 nl containing 80-200 pg of sgRNA and 100 pg of Cas9 mRNA. At 48 hpf, ten embryos were analysed for mosaicism at targeted loci using high resolution melt analysis. Injected embryos were raised to adulthood and back-crossed to identify transmitted mutations in F1 progeny. Homozygotes for each gene were generated by in-cross of F2 heterozygotes. Primers sequences used for genotyping by Sanger sequencing can be found as Supplementary Table S1 online.

### Quantitative real-time PCR

Total RNA was isolated from a pool of 20 mechanically homogenized embryos at 96 hpf for each sample (n=4) using TRItidy G (AppliChem) and 500 ng of cDNA was synthesized using QuantiTect Reverse Transcription Kit (Qiagen) according to manufacturer’s protocol. Primer sequences for *desma* (covering both *desma-1* and *desma-2*), *desmb* and *actb1* can be found as Supplementary Table S1 online. qPCR was performed in triplicates using SensiFAST SYBR No-ROX Kit (Bioline) in Rotor-Gene 6000 (Corbett Life Science). *Desma* and *desmb* expression levels were normalized to *actb1* expression, mutant expression levels were calculated relative to wild-type using ΔΔCt method^35^.

### Protein Extraction and Immunoblotting

For protein extraction, 1-year-old fish were euthanized using overdose of tricaine (MS222, 300 mg/l). Skeletal muscle tissue was immediately dissected and fresh-frozen in liquid nitrogen. Tissue was pulverized and homogenized in blending buffer (16 M Tris HCl, 200 M EDTA, 20% SDS, 1X protease inhibitor cocktail, volume adjusted to 5 ml with dH_2_O) by sonication on ice. Lysates were then centrifuged and supernatant transferred in a fresh tube. Total protein concentrations were determined by bicinchoninic acid Protein Assay (Pierce, 23225). Equal amounts of protein (40 µg) were loaded on a 13% polyacrylamide gel and transferred onto a nitrocellulose membrane using semi-dry transfer, blocked for one hour at room temperature (%5 nonfat dried milk in %0,2 TBS-T) and probed with anti-desmin antibody (1:1000; Sigma-Aldrich, D8281) overnight at 4°C or anti-lamin B1 antibody (Abcam, ab90169) followed by appropriate HRP-conjugated secondary antibody. Chemiluminescence detection was done by using SuperSignal^™^ West Femto Maximum Sensitivity substrate (Thermo Scientific, 34095) in GeneGnome device. See Supplementary Fig. S5 for uncropped full-length pictures of western blotting membranes.

### Whole-mount Immunofluorescence

96 hpf embryos were fixed in %4 PFA overnight at +4°C and washed with PBTx (%0.01 Triton X-100 in PBS). Embryos were permeabilized in 50 µg/ml Proteinase K for 2 hours and re-fixed in 2% PFA for 20 minutes at room temperature. Embryos were blocked in %5 BSA for 2 h at room temperature and incubated with anti-desmin antibody (1:20; Sigma, D8281) for two overnights. After two overnight incubations with Alexa-Fluor 488 conjugated secondary antibody, embryos were mounted with DAPI onto glass bottom dishes for imaging with Zeiss LSM Pascal laser scanning confocal microscope.

### Larval Phenotype and Morphology

After successful mating, embryos were collected, counted and recorded for the following rates. Hatching rate was reported as cumulative percentage of hatched embryos at 24 hpf, 48 hpf and 72 hpf. Mortality rate was expressed as the percentage of death embryos for 5 dpf. For body length measurements, at least 16 embryos at 96 hpf were mounted, photographed (The Imaging Source, DFK 41AU02) and measured from the mouth tip to the tail base using ImageJ (Version 1.49u). Values for body length were presented as mean body length in cm. All the experiments except body length measurements (three times) were performed at least eight times.

### Phalloidin Staining

Phalloidin staining was performed on 96 hpf embryos (N=8) as described^36^. Following fixation in 4% PFA and permeabilization with 50 µg/ml Proteinase K, embryos were incubated in Alexa-Fluor 488 conjugated Phalloidin (Invitrogen, A12379) overnight at +4°C. Embryos were then mounted onto glass bottom dishes and photographed with Zeiss LSM Pascal laser scanning confocal microscope. Somite lengths were measured by using ImageJ (Version 1.49u).

### Hematoxylin Eosin Staining

Adult zebrafish were freshly frozen in methyl-butanol cooled down in liquid nitrogen and then taken into O.C.T compound for sectioning. 8 µm-thick tissue sections were incubated in hematoxylin for 5 minutes, 1% acid-alcohol solution for 1 minute, 1% ammonia for 1 minute, eosin solution for 3 minutes respectively by washing with distilled water in between steps. Finally, sections were washed with alcohol and xylene, and mounted. Sections were examined under Nikon Eclipse E-400 microscope and photographed by DXM 1200F digital camera.

### Transmission Electron Microscopy

Fixation was performed by immersion in 2% glutaraldehyde in PBS for 3 hours, and the post fixation by 1% osmium tetroxide in the dark for 1 hour at room temperature. Fixed tissues were dehydrated in graded ethanol series, cleared in propylene oxide and embedded in epon (Agar Scientific, UK). Forty two-sixty nanometer thick ultrathin sections were stained with uranyl acetate and lead citrate, analyzed under a transmission electron microscope (JEOL-JEM 1400, Japan) and attached CCD camera (Gatan Inc., Pleasanton, CA, USA)^37^. Striated muscle ultrastructure has been evaluated for sarcolemma, myofibrils and cytoskeleton, nucleus, mitochondria, membrane systems, deposits and particles and other unusual structures^38^.

### Motility Assay

For motility assay 48 hpf embryos from in-crosses of *desma*^*kg97/+*^, *desmb*^*kg156/+*^ or *desma*^*kg97/+*^;*desmb*^*kg156/kg156*^ adults were used. Motility assays were performed by previously described protocol^18^. A petri dish was placed on a transparent sheet with concentric circles 5 mm apart. One embryo at a time was positioned in the center of the dish and stimulated using a dissection needle. The escape response of the embryo was recorded using a high-speed camera (Huawei, Mate 10 Lite, 16 MP, 120 fps). The movements of the embryo—midpoint between the two eyes—were tracked using a custom software that works based on template matching algorithm. The elapsed time between the last frame before touch stimulus and the first frame after the body of the fish contacts the 10 mm circle was computed from the digitized embryo movements with a resolution of 8.3 ms (120 fps). Experiments were performed blindly and after video accusation, DNA was extracted from each embryo for genotyping.

### Calcium Flux

1-year-old fish were euthanized using overdose of tricaine (MS222, 300 mg/l). After removal of the skin, tissue samples were incubated into 5 mg/ml collagenase solution in PBS (without Ca^2+^ and Mg^2+^) at 28.5 °C for 20 minutes on a shaker at 50 rpm. Samples have been triturated gently under dissecting microscope in order to dissociate muscle fibres from the tissue components until rode-shaped muscle fibres were obtained. Dissociated fibres were incubated in DMEM containing 10% FBS and 1% antibiotic-antimycotic overnight at 28.5 °C and 5% CO_2_ environment. On the following day, fibres were incubated in 2.5 μM Fluo-4 AM for 20 minutes in 5% CO_2_. Under a laser scanning confocal microscope (Zeiss LSM Pascal), glass microelectrodes (filled with 3M KCl) were inserted into muscle fibres in a current clamp mode^39^. A depolarizing current stimulus was delivered through the microelectrode to excite the fibre and evoke a calcium signal. The evoked calcium-specific fluorescent emission signal was recorded in “Line Scan” mode of confocal microscope, supplying information about a line cross sectioning the muscle fibre at a high frequency in tandem with the electrical recordings. Data were analyzed in MATLAB (The MathWorks, Inc). The stimulus waveform, which consisted of four consecutive pulses increasing in duration, was used in all the experiments. The amplitude of the current pulse was kept constant at 100 nA while the durations were 10 ms, 20 ms, 30 ms, 40 ms respectively. Experiments were done by using three WT, three *desma*^*kg97/kg97*^ and two *desmb*^*kg156/kg156*^ fish and at least eight fibres from each group were recorded.

### Statistics

Data were statistically analyzed by GraphPad Prism 8 (GraphPad Software Inc.) by nonparametric Mann-Whitney U test (Two-tailed) when two groups were compared. The results were considered significant when *P*<0.05. For analysis of time course experiments (hatching rate, mortality rate and time course calcium flux), repeated measures two-way ANOVA was used and Bonferroni post *hoc* test was used for multiple comparisons. All error bars were presented as mean±SEM and bars were represented as median.

## Acknowledgements

This work was supported by The Scientific and Technological Research Council of Turkey, Project no. 214S174 to P.R.D. S.M.H. is an MRC Scientist with MRC Programme Grant G1001029 and MR/N021231/1 support.

## Competing interests

The author(s) declare no competing interests.

## Data Availability

Supporting information is available in Supplementary files online and further information is available from the corresponding author upon request.

## Author Contributions

P.R.D. designed, P.R.D. and S.M.H. supervised the study. G.K.K. wrote the paper and assembled the figures, E.K.M. made contribution to writing the paper. G.K.K., E.K.M, C.K., S.U., B.S., B.E., M.G., I.U., N.B.D., P.K., B.T. and N.P. performed the experiments and/or analyzed the results. All authors interpreted the data and edited the manuscript.

## Notes

### Competing Interest Statement

The authors have declared no competing interest.

