## Supplementary Figures for "Knockout of zebrafish desmin genes does not cause skeletal muscle degeneration but alters calcium flux"

### INVENTORY OF SUPPLEMENTARY INFORMATION

#### 1. SUPPLEMENTARY FIGURES AND LEGENDS

**Figure S1:** *Desma* and *desmb* gene and transcript structures.

**Figure S2:** ISH with sense *desma* and *desmb* probes.

**Figure S3:** Quantification of somite length, number of nuclei and fibre diameter.

**Figure S4:** Relative *desma, desmb* and *vim* mRNA expression levels.

**Figure S5**: Uncropped pictures of western blots.

#### 2. SUPPLEMENTARY METHOD

*In silico* prediction of transcription factors binding to *desma* and *desmb* promoters.

#### 1. SUPPLEMENTARY FIGURES AND LEGENDS

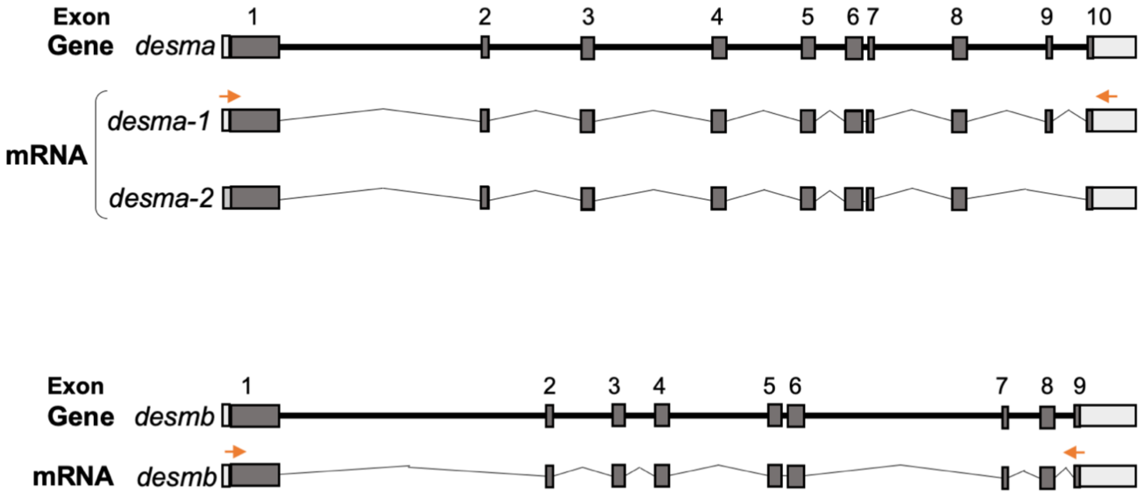


**Figure S1.** Schematic drawings of *desma* gene and mRNA transcripts, and *desmb* gene and mRNA transcript. Exons are represented as dark grey boxes, UTRs as light grey boxes and introns as connecting lines. Corresponding location of primers used for synthesizing antisense probes for in situ hybridization are shown as orange arrows.

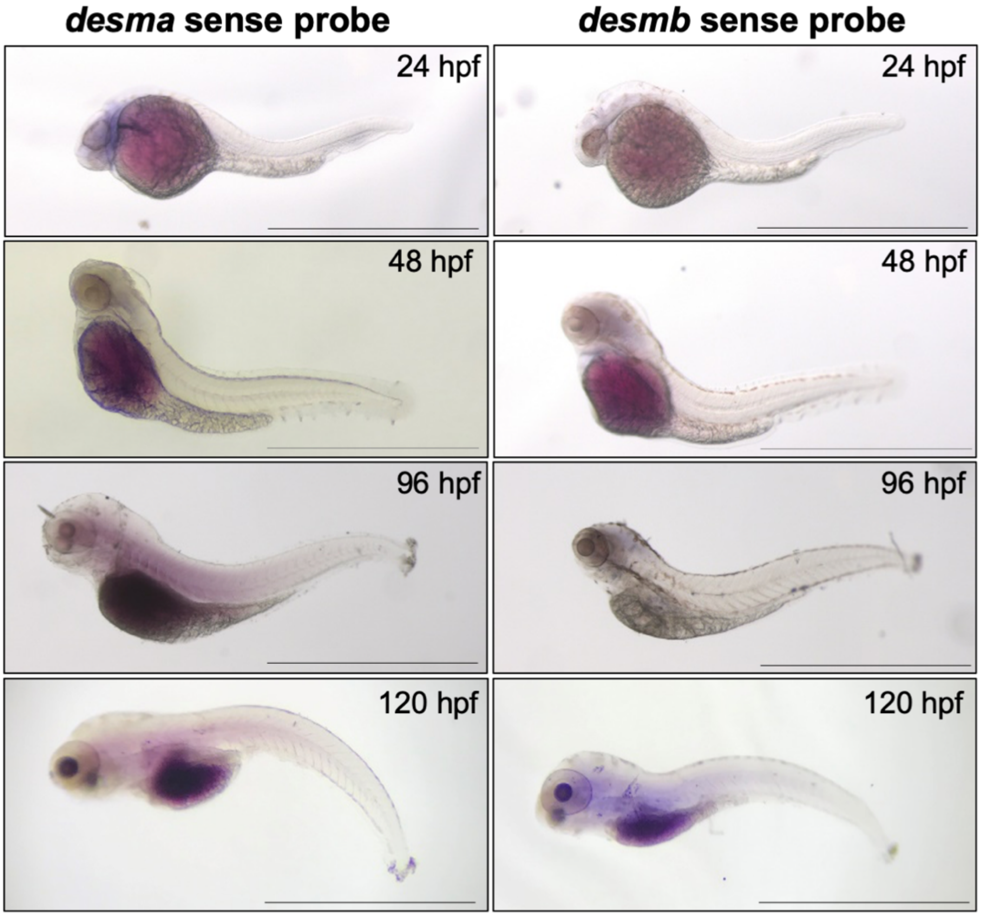


**Figure S2.** Whole mount in situ mRNA hybridisation of WT embryos at the indicated stages for sense probes to *desma* or *desmb.* Scale bar: 1 mm.

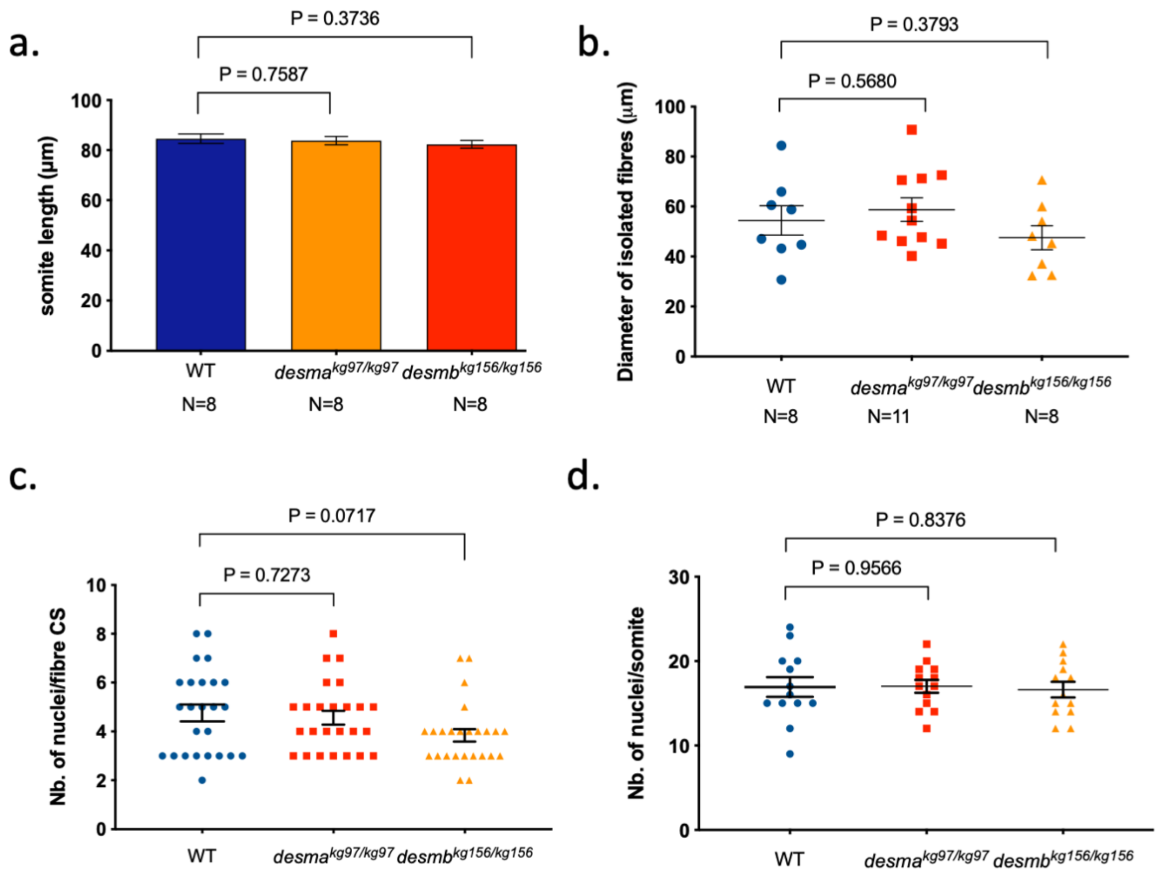


**Figure S3.** (**a)** Somite lengths (µm) were measured from optical sections of WT and mutant embryos stained with rhodamine phalloidin (Fig. 4e) (N=8, Mann-Whitney U). (**b**) Diameter of isolated skeletal muscle fibres from 1-year old adult skeletal muscle tissue (µm,) used in calcium flux assay (N>=8, Student's t-test). (**c**) Number of nuclei per fibre cross-section (CS) were calculated from hematoxylin-eosin staining of 1-year old adult skeletal muscle tissues (N=25, Student's t-test). (**d**) Number of nuclei per somite was calculated by counting the number of nuclei in optical sections of the mid-trunk of 96 hpf embryos stained with DAPI by confocal microscopy (N=13, Student's t-test).

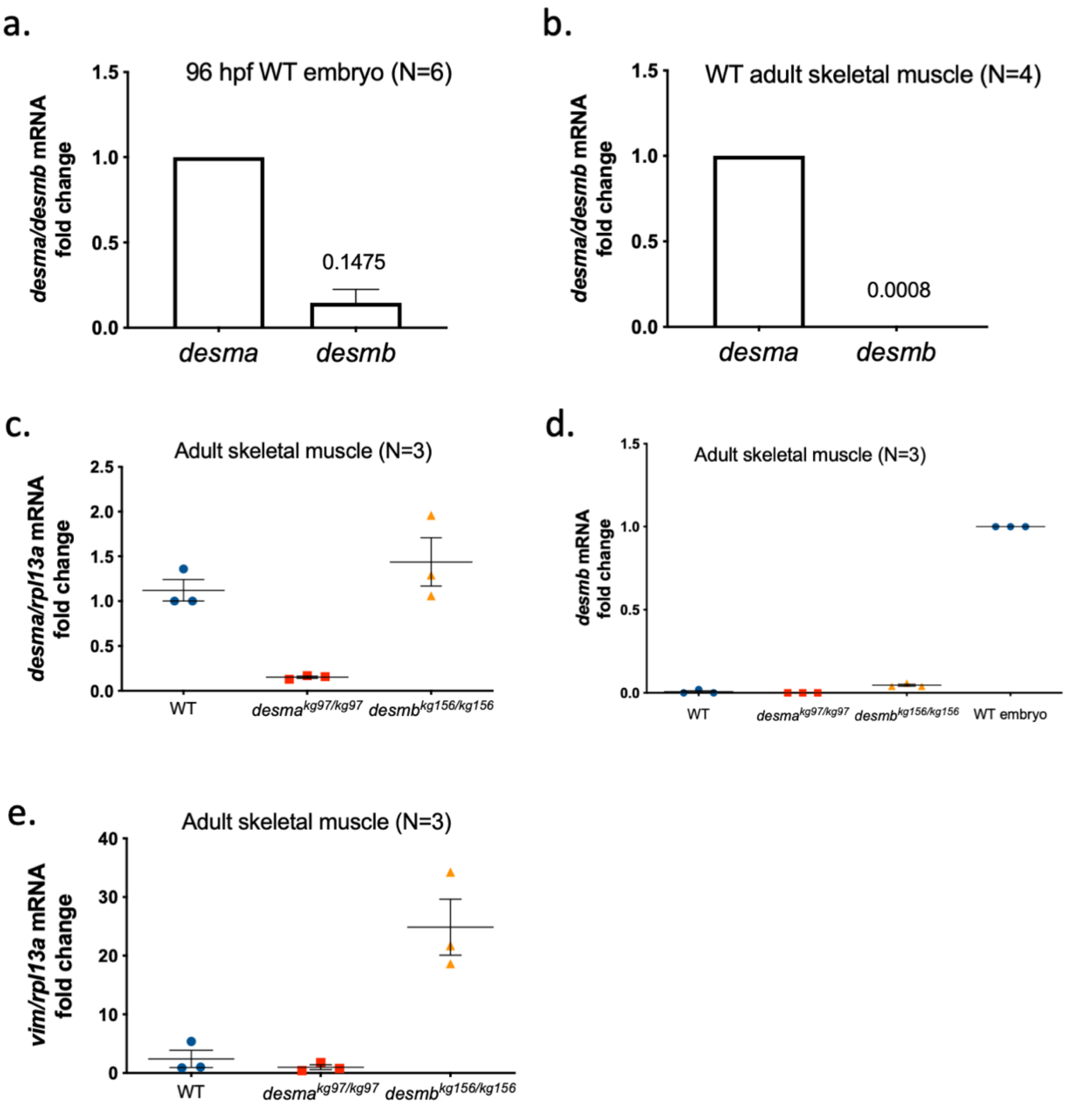


**Figure S4. (a)** Relative *desma* and *desmb* mRNA expression levels in WT 96 hpf embryos (N=6). *Desma* and *desmb* Ct values were normalized to *rpl13a* and *actb1*. (**b**) Relative *desma* and *desmb* mRNA expression levels in WT adult skeletal muscle tissue samples (N=4). *Desma* and *desmb* Ct values were normalized to *rpl13a*. (**c**) *Desma* mRNA fold change in WT and mutant adult skeletal muscle tissue samples (N=3). Expression was normalized to *rpl13a*. (**d**) *Desmb* mRNA fold change in WT and mutant adult skeletal muscle tissue samples, *desmb* mRNA level in 96 hpf WT embryo has been set to 1 (N=3). Expression was normalized to *rpl13a*. (**e**) *Vimentin* mRNA fold change in WT and mutant adult skeletal muscle tissue samples (N=3). Expression was normalized to *rpl13a*.

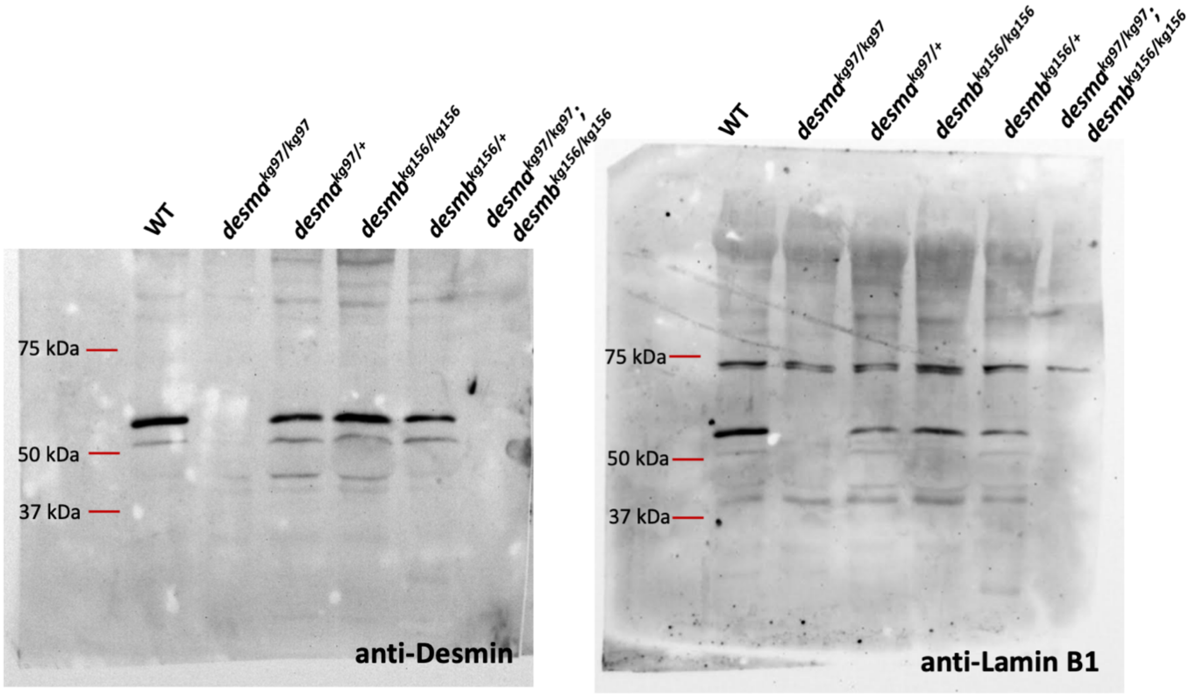


**Figure S5.** Uncropped full-length pictures of the western blotting membrane presented in Fig. 4a. Left picture shows membrane incubated with anti-desmin antibody (Sigma-Aldrich, D8281) followed by HRP-conjugated secondary antibody and (90 sec. exposure). Right picture shows the same membrane incubated with anti-lamin B1 antibody (Abcam, ab90169) after several washes, followed by HRP-conjugated secondary antibody (120 sec. exposure).

#### 2. SUPPLEMENTARY METHOD

##### *In silico* prediction of transcription factors binding on *desma* and *desmb* promoters

First, The Eukaryotic Promoter Database (EPD)[1] was used to retrieve promoter sequences for *desma* (ENSDARG00000058656) and *desmb* (ENSDARG00000005221) genes. While a defined core promoter was annotated for *desma*, no information was available for *desmb*. Therewith, sequences up to 2,000 bp upstream of transcription start sites for *desma* and *desmb* were retrieved from Ensembl and analyzed for prediction of transcription factor (TF) binding sites on AnimalTFDB 3.0[2]. Lists obtained for each gene were matched and TFs common to *desma* and *desmb*, or unique to *desma* and unique to *desmb* were identified. Lastly, transcription factors were annotated using Enrichr tool[3].


1. Dreos, R., Ambrosini, G., Groux, R., Perier, R. C. & Bucher, P. The eukaryotic promoter database in its 30th year: Focus on non-vertebrate organisms. *Nucleic Acids Res.* **45**, D51–D55 (2017).
2. Hu, H. *et al.* AnimalTFDB 3.0: A comprehensive resource for annotation and prediction of animal transcription factors. *Nucleic Acids Res.* **47**, D33–D38 (2019).
3. Chen, E. Y. *et al.* Enrichr: Interactive and collaborative HTML5 gene list enrichment analysis tool. *BMC Bioinformatics* (2013) doi:10.1186/1471-2105-14-128.
