## Supplementary Table S1 for "Knockout of zebrafish desmin genes does not cause skeletal muscle degeneration but alters calcium flux"

**Table S1.** List of primer sequences used in for synthesis of ISH probes, genotyping and qPCR experiments.

| | **Forward primer sequence** | **Reverse primer sequence** |
|---|---|---|
| *desma* antisense probe | 5'-TACATCGAGAAGGTGCGCTT-3' | 5'-GGATCCATTAACCCTCACTAA AGGGAATTGTCTCCATGCGTCAT CCA-3' |
| *desmb* antisense probe | 5'-AATGACCGCTTCGCCAACTA-3' | 5'-GGATCCATTAACCCTCACTAA AGGGAACCTCTCCATCACGTGTC TCG-3' |
| *desma* sense probe | 5'-TAATACGACTCACTATAGGGA GATACATCGAGAAGGTGCGCTT-3' | 5'-TTGTCTCCATGCGTCATCCA-3' |
| *desmb* sense probe | 5'-TAATACGACTCACTATAGGGAG AAATGACCGCTTCGCCAACTA-3' | 5'-CCTCTCCATCACGTGTCTCG-3' |
| *desma* genotyping primers | 5’-CGCACGGTGGCTGGAAAA-3’ | 5’-CACGGTTCCTCAGGTTCCTC-3’ |
| *desmb* genotyping primers | 5’-GTCATCCACTTCTTCTCCTGG-3’ | 5’-GTGCGAGTGTGGAGAAACTC3’ |
| *desma* qPCR primers | 5’-GCTGCCAAGAATATCAGCGA-3’ | 5’-TGCCTCCTCAGAGACTCATT-3’ |
| *desmb* qPCR primers | 5’-ATGCAAGAGACCCAAGTCCA-3’ | 5’-TCTTGCTCACAGCCTGGTTA-3’ |
| *actb1* qPCR primers | 5’-TCTTCCAGCCTTCCTTCCTG-3' | 5’-TGTGTTGGCATACAGGTCCT-3’ |
| *vim* qPCR primers | 5’-ACCTAGGGGAGGACATCGAG-3’ | 5’-CGAGCCAGAGAGGCGTTATC-3’ |
| *rpl13a* qPCR primers | 5'-TCTGGAGGACTGTAAGAGGTATGC-3' | 5'-AGACGCACAATCTTGAGAGCAG-3' |
